# From sequence to scaffold: computational design of protein nanoparticle vaccines from AlphaFold2-predicted building blocks

**DOI:** 10.1101/2025.08.20.671178

**Authors:** Cyrus M. Haas, Naveen Jasti, Annie Dosey, Joel D. Allen, Rebecca Gillespie, Jackson McGowan, Elizabeth M. Leaf, Max Crispin, Cole A. DeForest, Masaru Kanekiyo, Neil P. King

**Affiliations:** Department of Chemical Engineering, University of Washington, Seattle, WA 98195, USA; Institute for Protein Design, University of Washington, Seattle, WA 98195, USA; Institute for Stem Cell and Regenerative Medicine, University of Washington, Seattle, WA 98195; Department of Molecular Engineering, University of Washington, Seattle, WA 98195, USA; Department of Biochemistry, University of Washington, Seattle, WA 98195, USA; School of Biological Sciences, University of Southampton, Southampton SO17 1BJ, UK; Vaccine Research Center, National Institute of Allergy and Infectious Diseases, National Institutes of Health, Bethesda, MD, 20892; Department of Bioengineering, University of Washington, Seattle, WA 98195, USA; Department of Chemistry, University of Washington, Seattle, WA 98195, USA; Molecular Engineering & Sciences Institute, University of Washington, Seattle, WA 98195, USA

**Keywords:** protein design, nanoparticles, vaccines, influenza

## Abstract

Self-assembling protein nanoparticles are being increasingly utilized in the design of next-generation vaccines due to their ability to induce antibody responses of superior magnitude, breadth, and durability. Computational protein design offers a route to novel nanoparticle scaffolds with structural and biochemical features tailored to specific vaccine applications. Although strategies for designing new self-assembling proteins have been established, the recent development of powerful machine learning-based tools for protein structure prediction and design provides an opportunity to overcome several of their limitations. Here, we leveraged these tools to develop a generalizable method for designing novel self-assembling proteins starting from AlphaFold2 predictions of oligomeric protein building blocks. We used the method to generate six new 60-subunit protein nanoparticles with icosahedral symmetry, and single-particle cryo-electron microscopy reconstructions of three of them revealed that they were designed with atomic-level accuracy. To transform one of these nanoparticles into a functional immunogen, we reoriented its termini through circular permutation, added a genetically encoded oligomannose-type glycan, and displayed a stabilized trimeric variant of the influenza hemagglutinin receptor binding domain through a rigid *de novo* linker. The resultant immunogen elicited potent receptor-blocking and neutralizing antibody responses in mice. Our results demonstrate the practical utility of machine learning-based protein modeling tools in the design of nanoparticle vaccines. More broadly, by eliminating the requirement for experimentally determined structures of protein building blocks, our method dramatically expands the number of starting points available for designing new self-assembling proteins.

**Significance Statement:** Self-assembling protein nanoparticle vaccines have steadily gained traction in both academic and industry vaccine development over the last decade. Recent work has shown that computationally designing new self-assembling proteins allows the structural and functional features of nanoparticle vaccines to be precisely tailored, and that this can significantly affect the vaccine-elicited immune response. To date, designing such nanoparticle vaccines has required the use of known crystal structures as starting points. Here we show how new machine learning tools can be leveraged to design new self-assembling protein nanoparticle immunogens in the absence of experimentally determined structures of the building blocks that elicit strong immune responses in mice.

## Introduction

The design of nanoparticle immunogens based on self-assembling protein scaffolds is a burgeoning area of technological innovation in vaccines (1–4). It has long been known that repetition of epitopes in three-dimensional space is a danger signal sensed by B cells through pathogen- or vaccine-driven clustering of their antigen receptors (5, 6). More recently, engineered protein nanoparticle immunogens and DNA nanotechnology objects have been used to explore in detail the effects of repetitive antigen display on B cell responses, highlighting the role of the detailed geometry of antigen presentation (7–13). In parallel, the advantageous trafficking of particulate immunogens *in vivo* and how they are sensed by the innate immune system is also becoming increasingly clear (14, 15). Together, these observations have motivated many efforts over the last decade in which antigens of interest are displayed on self-assembling protein scaffolds, beginning with two landmark studies in 2013 (16, 17). These studies generally show that protein nanoparticle immunogens elicit significantly more potent humoral immune responses than soluble antigen, and in some cases have also demonstrated improved quality, breadth, and durability (9, 18–20). Many of these efforts have used naturally occurring self-assembling proteins such as ferritin and lumazine synthase as scaffolds (16, 17), although computationally designed protein nanoparticles are being increasingly adopted by the field (21–23). Importantly, a COVID-19 vaccine comprising the receptor binding domain (RBD) of the SARS-CoV-2 spike protein displayed on a computationally designed protein nanoparticle was recently licensed in multiple countries, clinically de-risking this technology (24–26).

All of the computationally designed protein nanoparticle vaccines studied to date have used protein nanoparticles generated by docking oligomeric building blocks in a target symmetric architecture and designing non-covalent protein-protein interfaces that drive assembly (27–29). Since its introduction more than a decade ago, this method has been generalized to design a large number of self-assembling proteins with a wide variety of structural properties (30–32). Nevertheless, except for a small number of custom oligomeric building blocks (21), this approach has otherwise been limited by a requirement for experimentally solved structures used as inputs to docking. Despite the large and growing Protein Data Bank (PDB; (33)), this remains a significant limitation, as available structures represent only a tiny fraction of the possible space of protein structures (34–36). For example, if ten unique trimers from the PDB each yield five promising nanoparticle designs, the result would be 50 possible nanoparticles. However, if the number of building block candidates could be expanded to 200 by leveraging homologous sequences, the result would be 1,000 possible nanoparticles, a 20-fold increase. Even within protein families that are represented in the PDB, biotechnologically relevant properties such as stability, secretability, and post-translational modifications can vary widely among homologs from different species (31, 37). Given that the structural and biophysical properties of an assembled nanoparticle are largely determined by the properties of its building blocks, methods that expand the number of protein building blocks available for design would facilitate the design of novel nanoparticles and nanoparticle vaccines with desired properties.

New machine learning (ML)-based methods are revolutionizing protein structure prediction and design (38–43). Design methods like RFdiffusion for backbone generation and ProteinMPNN for amino acid sequence design have dramatically increased the success rate of many *de novo* design challenges, aided by the use of structure prediction methods such as AlphaFold2 (AF2) and RoseTTAFold as filters for high-quality designed proteins (44–46). For example, in protein nanoparticle design specifically, two recent papers highlighted that using ProteinMPNN for interface design resulted in improved interfaces and nanoparticles compared to traditional Rosetta-based sequence design (47, 48). Although the ability of AF2 and RoseTTAFold to provide accurate models of protein structures from sequence alone provides an opportunity to dramatically expand the number of building blocks available for nanoparticle design, no novel self-assembling nanoparticles based on predicted structures from these tools have been reported yet.

Here, we develop a new and general method for designing self-assembling proteins that uses predicted structures of oligomeric proteins from thermophilic organisms as building blocks. We found that novel homomeric nanoparticles can be designed with atomic-level accuracy from sequence alone, and can be engineered to elicit robust antibody responses against influenza by repetitively displaying antigens based on the RBD of the viral hemagglutinin.

## Results

### Computational nanoparticle design

We set out to leverage ML-based structure prediction tools to expand the number of building blocks available for novel nanoparticle design. Our general strategy was to i) predict the structures of homologs of oligomeric proteins of known structure, and ii) use the predictions as inputs to established pipelines for nanoparticle design based on symmetric docking and protein-protein interface design (49). We focused on the design of homomeric (“one-component”) nanoparticles, as the docking of such assemblies is severely constrained and would benefit most from the availability of a wider variety of building blocks. Based on our prior experiences designing one-component nanoparticles (28, 31, 49), we attempted to identify building blocks with attributes that may increase the likelihood they would assemble into nanoparticles. Specifically, we searched the PDB for homomeric proteins from bacteria or archaea that had 3-fold rotational symmetry (C3), high-resolution structures (≤2.4 Å), and significant (>50%) helical secondary structure (**Fig. 1A**). After clustering similar sequences, 95 structures (PDB seeds) passed these filters. We then screened the UniProt database for proteins from thermophilic organisms with >50% sequence identity to these 95 PDB seeds. Previous work has shown thermophilic homologs can be up to 10-fold more tolerant of mutations at individual residues than their mesophilic counterparts, allowing for the evolution of thermostable proteins into binders, receptor analogs, and P450 enzymes with novel activities (50–56). By focusing on proteins from thermophiles, we hoped to identify starting structures that were robust, soluble, and tolerant to mutation. Of the 359 identified thermophilic trimer homologs, 191 yielded high-confidence AF2 predictions (pAE < 5, pLDDT > 90) (**Fig. 1B**). We docked these predicted structures in the so-called “I3” architecture (28) by aligning their three-fold symmetry axes to the corresponding axes of icosahedral point group symmetry, using the Rosetta SymDofMover to sample the translational and rotational degrees of freedom along this shared axis (**Fig. 1C**, (27)). We then designed the interfaces between the docked C3 building blocks using ProteinMPNN (**Fig. 1D and *SI Appendix*, Table S1**), which was recently shown to generate high-quality nanoparticle interfaces without requiring manual intervention ((47, 48); *Materials and Methods*). This pipeline produced ∼430,000 designs, which were filtered using Rosetta scoring metrics (ddG < −10, buried Solvent Accessible Surface Area (SASA) > 300, Shape Complementarity > 0.5, and dGins,pred > 2.6; ***SI Appendix*, Fig. S1**) and by visual inspection to select 88 designs for experimental characterization.

**Fig. 1.**
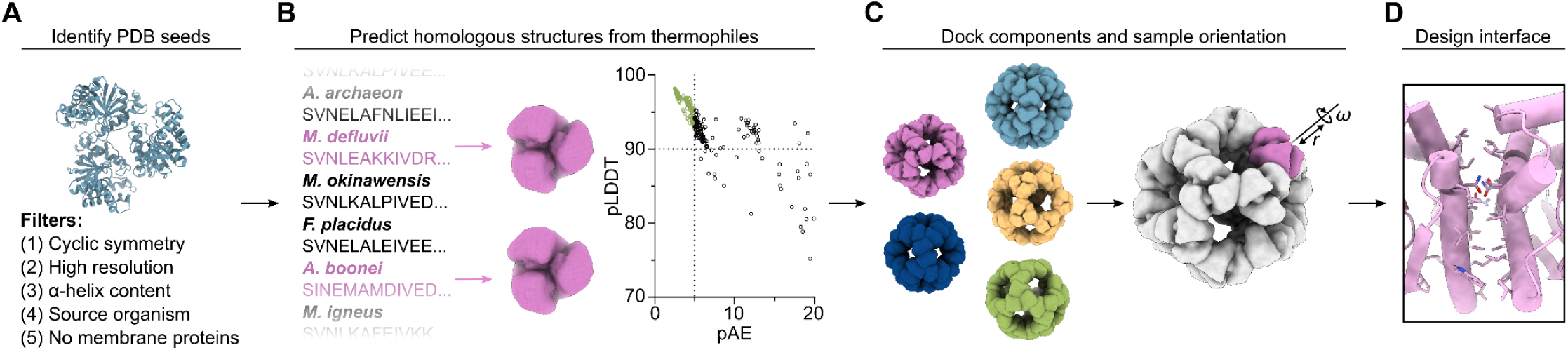
Computational design of novel self-assembling proteins from AF2-predicted models. *(A)* The PDB was searched for structures that possess optimal attributes to form nanoparticles. *(B)* From these PDB seeds, homologous sequences were identified from thermophilic organisms and their structures predicted with AlphaFold2. *(C)* The best-scoring predictions were docked using RPXDock (40) and their translational (r) and rotational (ω) degrees of freedom were subsequently sampled using the Rosetta SymDofMover (27) to generate multiple structures for interface design. *(D)* The interfaces between docked nanoparticle components were designed using ProteinMPNN (42) and a subset of these designs was selected using Rosetta scoring metrics for experimental characterization.

### Screening and characterization of designed nanoparticles

We ordered genes for expression in *Escherichia coli* encoding the 88 designs with hexahistidine tags appended to enable purification via immobilized metal affinity chromatography (IMAC). IMAC-purified samples were subsequently analyzed by size-exclusion chromatography (SEC), resulting in ten putative hits with elution peaks at the expected retention time (**Fig. 2A**). This initial success rate compares favorably to historical success rates for established design protocols starting from building blocks of known structure. Following large-scale expression and purification of these hits, SEC confirmed that six designs yielded predominant peaks at the expected elution volume, with three of the six also featuring substantial peaks at the void volume of the column (**Fig. 2B**). These six designs originated from three unique PDB seeds (6HS8: I3-A6, I3-B2, I3-B3, and I3-B5; 1QWG: I3-D12; 6HF7: I3-A7) and therefore have three distinct backbones (***SI Appendix*, Table S2**). SEC fractions at the expected elution volume were pooled and analyzed by dynamic light scattering (DLS) and negative stain electron microscopy (nsEM), which revealed the presence of homogeneous nanoparticles for all six samples. Low-resolution 3D reconstructions from nsEM were consistent with the computational design models. We measured the aggregation temperature (*T_agg_*) for each nanoparticle by monitoring DLS while heating from 25 to 84°C, which showed that all nanoparticles retained their intended assembly state until at least 60°C, and I3-D12 did not aggregate in the measured range (**Fig. 2C and *SI Appendix*, Fig. S2**).

**Fig. 2.**
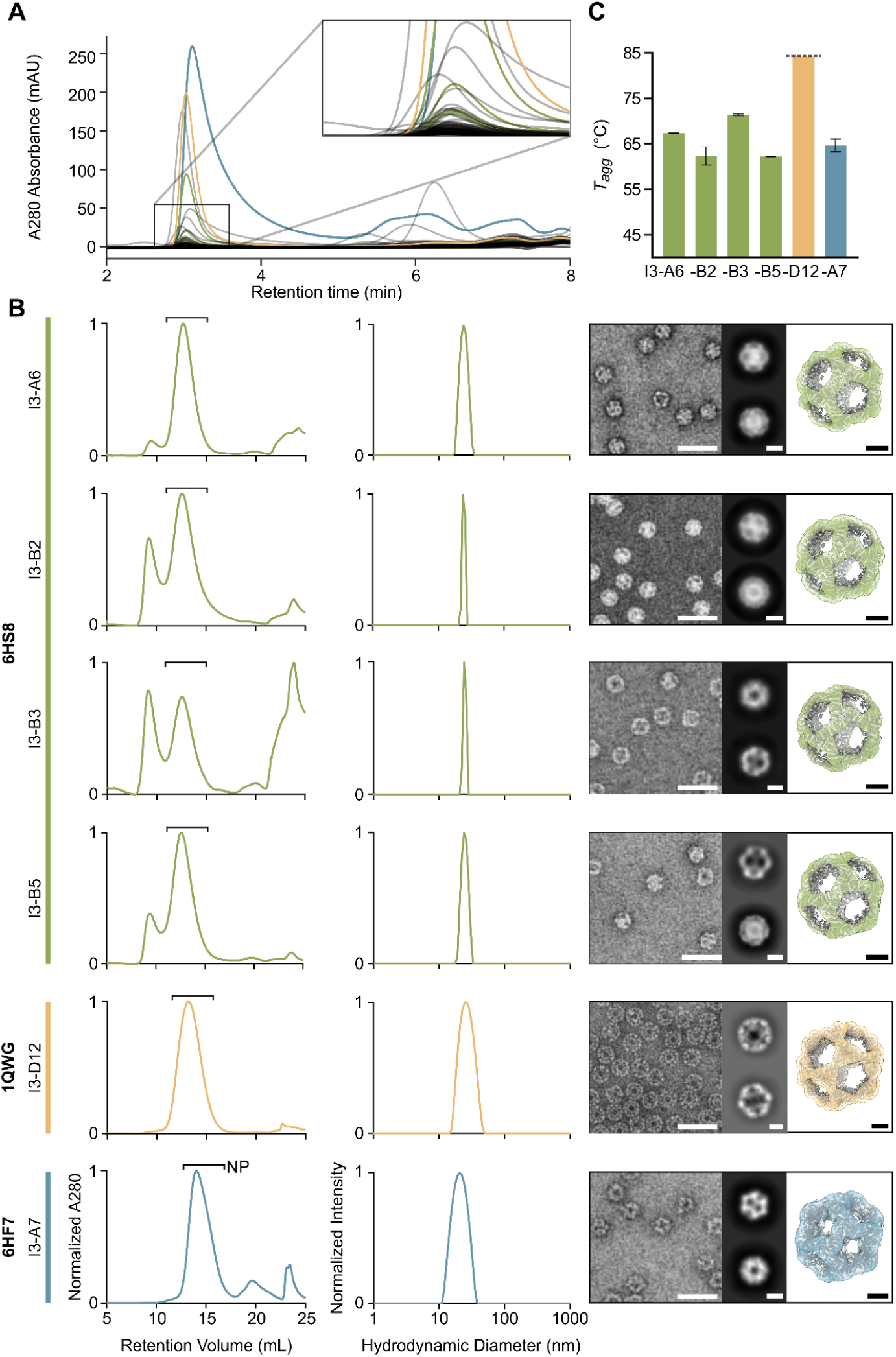
Screening and characterization of designed nanoparticles. *(A)* An overlay of HPLC traces for all 88 designs tested at small scale. The magnified region shows a large number of putative hits at the expected retention time for icosahedral nanoparticles. *(B)* From left to right, normalized SEC traces of IMAC eluent from large-scale expression, normalized DLS, and negative stain micrographs with 2D averages and 3D reconstructions for each of the six confirmed nanoparticles. Bars labeled “NP” indicate the SEC fractions containing nanoparticles that were pooled for further characterization. Scale bars represent 40 nm, 10 nm, and 5 nm for the micrographs, 2D averages, and 3D reconstructions, respectively. *(C)* Aggregation temperatures for the six confirmed nanoparticles. Data related to nanoparticles from PDB seeds 6HS8, 1QWG, and 6HF7 are colored in blue, yellow, and green, respectively.

### High-resolution structural characterization of three nanoparticles

To evaluate the accuracy of our design method at high resolution, we used single-particle cryo-electron microscopy (cryo-EM) to determine the structures of one nanoparticle from each of the successful PDB seeds. Homogeneous and monodisperse particles were observed in the raw micrographs for all three samples, and 2D class averages exhibited multiple well-defined views that provided clear evidence of icosahedral symmetry (**Fig. 3A**). Icosahedral 3D reconstructions at 2.5 Å, 2.3 Å, and 3.5 Å resolution were obtained for I3-A6, I3-D12, and I3-A7, respectively, allowing us to build atomic models of each nanoparticle (**Fig. 3B; *SI Appendix*, Fig. S3 and Table S3**). Comparing the experimental cryo-EM structures to the computational design models demonstrated that AF2 structure prediction was quite accurate: the all-atom root mean square deviations (RMSDs) for a single monomer of I3-A6, I3-D12 and I3-A7 were 1.3 Å, 1.7 Å, and 1.4 Å, respectively. Aligning two subunits across the designed interfaces revealed that these were designed with high accuracy, yielding alpha-carbon RMSDs of 0.7 Å, 1.1 Å, and 1.5 Å, respectively (**Fig. 3C,D**). In each case, the rotameric states of the interdigitated hydrophobic side chains at the designed interfaces corresponded well with the design models, as did the polar residues lining the periphery of the I3-A6 and I3-D12 interfaces (**Fig. 3D**). Several of the polar residues at the I3-A7 interface adopted unintended rotamers, likely due to small shifts in the backbones of the loops that make up the interface. Nevertheless, essentially all of the polar and non-polar contacts at each interface recapitulated those in the design models. Slight rigid body deviations between the building blocks added up to alpha-carbon RMSDs of 1.6 Å, 2.2 Å, and 2.2 Å across the entire 60-subunit nanoparticle for I3-A6, I3-D12 and I3-A7, respectively. These values are comparable to those obtained from previous computational nanoparticle design campaigns that started from high-resolution crystal structures of the building blocks (27, 29, 47). These results establish that ML-based tools for structure prediction (AF2) and design (ProteinMPNN) enable the accurate design of novel self-assembling protein nanomaterials from predicted structures.

**Fig. 3.**
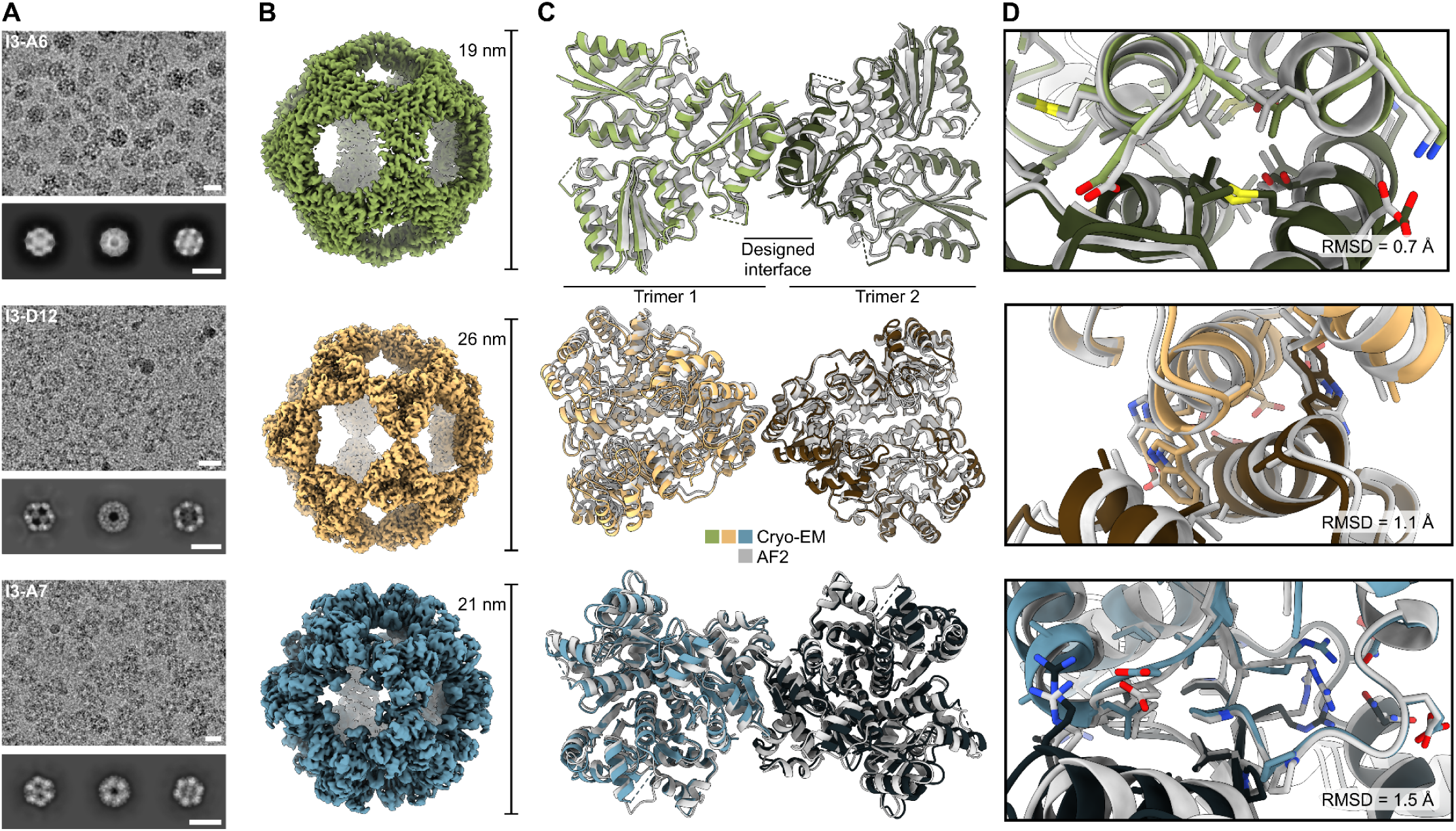
High-resolution structural characterization of three nanoparticles. *(A)* Processed micrographs and 2D class averages. Scale bars represent the diameter of each nanoparticle as labeled in panel B. *(B)* Cryo-EM reconstructions. *(C)* Two overlaid trimers across the two-fold axis. Dashed lines represent residues that were not modeled due to insufficient density. *(D)* Close-up of the designed interface for each nanoparticle. Gray structures represent AlphaFold2 predictions while colored ones are the true cryo-EM structure.

### Engineering I3-A7 to function as a nanoparticle immunogen

The application of computationally designed proteins as scaffolds for nanoparticle vaccines has advanced in recent years, with one such vaccine licensed (for SARS-CoV-2 (4, 24)) and several others in clinical trials for additional indications (8, 23). To test whether our designed assemblies could be functionalized as multivalent nanoparticle immunogens through genetic fusion of an antigen of interest, we set out to re-engineer several of their structural and biochemical properties. First, we evaluated whether they could be secreted in intact form from mammalian cells, a prerequisite for display of many glycoprotein antigens that require post-translational modifications found only in the secretory pathway (57). We found that only one of our six designed nanoparticles (I3-A7) successfully secreted from transfected Expi293F cells when an N-terminal signal peptide was appended, so our subsequent engineering efforts focused on this nanoparticle (***SI Appendix*, Fig. S4**). Second, as described below, we circularly permuted the subunit of I3-A7 so that its N terminus was present on the exterior of the nanoparticle and available for genetic fusion. Third, we engineered an oligomannose-type N-linked glycan into the I3-A7 subunit, as these have been shown to improve the trafficking and immunogenicity of particulate immunogens (3, 14). Finally, we designed a *de novo* rigid genetic linker between the re-engineered I3-A7 subunit and a variant of the influenza hemagglutinin (HA) RBD, stabilized in a native-like, closed trimeric conformation (“trihead”; (18)).

Display of the HA trihead via genetic fusion required the N terminus of I3-A7 to be accessible on the nanoparticle exterior. As the termini of I3-A7 were originally buried and therefore inaccessible, we circularly permuted the subunit, a genetic manipulation in which the original termini are fused together and new termini are created elsewhere (58). By visual inspection, we chose to “cut” the subunit between Asp133 and Ser134 (Genbank PKK86397 numbering), which would place the new N terminus on the exterior surface of each subunit (**Fig. 4A**). We then used RFdiffusion to generate *de novo* backbones for linkers connecting the two original termini (Gly2 and Asn183) up to four amino acids in length. The two adjacent amino acids on each side were allowed to move during diffusion in order to optimize backbone geometry throughout the linker. We then designed sequences for the new backbones using ProteinMPNN and predicted the structures of trimers of the circularly permuted subunits by AF2. We used visual inspection and AF2 prediction metrics for the diffused regions (pLDDT > 92, pAE < 5, IpTM > 0.9) to select four different linkers to test experimentally (**Fig. 4B**).

**Fig. 4.**
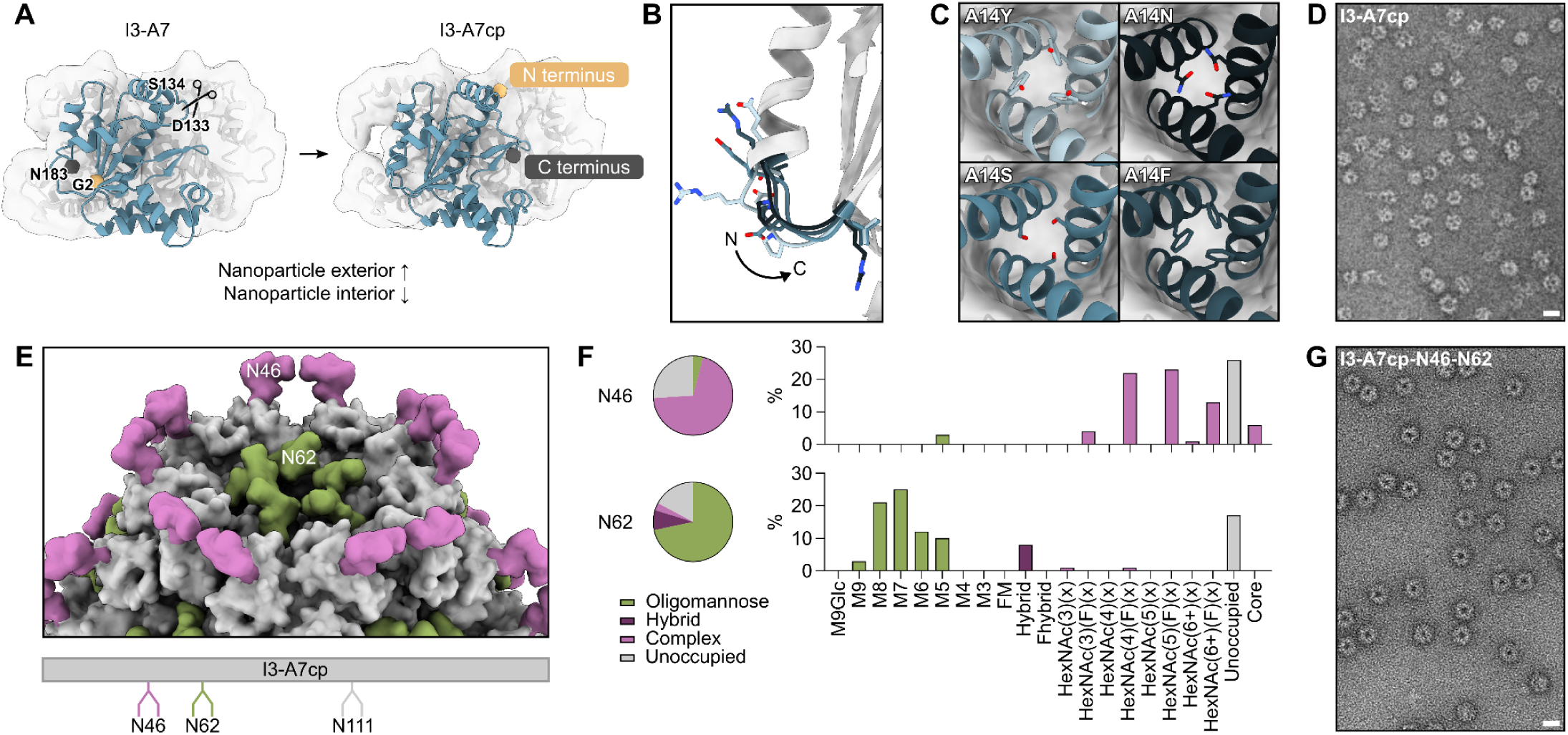
Engineering I3-A7 to function as a nanoparticle immunogen. *(A)* A schematic showing the process of circular permutation on a subunit of the trimer that forms the nanoparticle. *(B)* The four diffused linkers between the I3-A7 termini that were tested experimentally. The arrow indicates the directionality of the I3-A7cp backbone from the N to C terminus. *(C)* Four MPNN outputs at a central position in the intertrimer interface. *(D)* A representative electron micrograph of the circularly permuted nanoparticle. Scale bar = 21 nm. *(E)* A depiction of the two N-linked glycans added to each subunit trimer. Glycans were modeled as Man_5_GlcNAc_2_ using GlycoShape (63). An engineered sequon at position N111 was found to be unoccupied. *(F)* Complete characterization of each occupied glycosylation site via LC-MS. *(G)* A representative electron micrograph of the nanoparticle with glycans at positions N46 and N62. Scale bar = 21 nm.

Before validating the permutations, we noticed that our post-permutation termini were near the intratrimer interface of I3-A7, introduced by a new break in the polypeptide chain that could potentially destabilize the trimer. We therefore used ProteinMPNN to evaluate stabilizing mutations at ten positions at the trimer interface selected by inspection of the parent PDB seed (PDB ID 6HF7). After filtering on AF2 prediction metrics (pLDDT > 90, pAE < 8, IpTM > 0.9), we selected five sets of mutations in the central helix near the three-fold symmetry axis. For example, mutations at position Ala14 increased hydrophobic packing (A14Y, A14F) or introduced new hydrogen bonds (A14Y, A14S, A14N) between the trimer subunits (**Fig. 4C**). All 20 combinations of the five redesigned trimeric interfaces with the four diffused linkers had strong AF2 prediction metrics, including low RMSD to the original trimer prediction of the thermophilic homolog. We identified the highest-expressing variant from Expi239F cells (called I3-A7cp) by SDS-PAGE of cell culture supernatants and confirmed that it still formed the expected icosahedral nanoparticle using SEC and nsEM (**Fig. 4D and *SI Appendix*, Fig. S4**).

We then engineered N-linked glycans into I3-A7cp to further optimize its performance as a nanoparticle immunogen. Glycans have been shown to improve antigen secretion (59, 60), shield antibody responses against off-target epitopes (61), and—specifically in the case of oligomannose-type glycans—enhance vaccine trafficking and immunogenicity by engaging the innate immune system (3, 14). We introduced sequons encoding N-linked glycans at three solvent-exposed positions in I3-A7cp: N111, which is located on the edge of the subunit and points upward from within the pores of the assembled nanoparticle; N46, found at the apex of the subunit pointing out towards the exterior of the assembled nanoparticle; and N62, which is also located on the edge of the subunit but points into the pores of the assembled nanoparticle (**Fig. 4E**). Although N-linked glycans were not detectable at position 111 in nanoparticles purified from transfected cells, we observed 74% and 83% occupancy at N46 and N62, respectively (**Fig. 4F**). Glycosylation did not noticeably impact nanoparticle morphology: SEC, DLS, and nsEM of I3-A7cp-N46-N62 were essentially indistinguishable from I3-A7 and I3-A7cp (**Fig. 4G and *SI Appendix*, Fig. S4**). Furthermore, all three nanoparticles were highly thermostable, with *T_agg_* ≥ 72°C (***SI Appendix*, Fig. S4C**). LC-MS analysis revealed that while N46 was occupied by fully processed complex-type glycans, N62 comprised 71% underprocessed oligomannose-type glycans. Oligomannose-type glycans are typically processed in the endoplasmic reticulum (ER) and Golgi to become complex-type glycans, but this can be prevented by chemical or steric inhibition of the enzymes involved (62). Because no chemical inhibitors of glycan processing were added during transfection, this result suggested that glycan processing at N62 was sterically constrained. Based on the location of N62 in the nanoparticle, we suspected that this steric constraint may have resulted from nanoparticle assembly early in the secretory pathway, prior to initial processing of oligomannose-type glycan precursors in the ER. To probe this, we expressed and purified a trimeric variant of I3-A7cp containing the glycan sequons but lacking the computationally designed interface that drives nanoparticle assembly. Analysis of this protein confirmed that N62 was occupied almost exclusively by complex-type glycans, comprising only 4% oligomannose-type glycans (***SI Appendix*, Fig. S5**). These data demonstrate that we engineered a genetically encoded oligomannose-type glycan into I3-A7cp that depends on nanoparticle assembly early in the secretory pathway. We kept only the N62 glycan in the nanoparticle immunogens for influenza described below, reverting the sequons at positions 46 and 111.

### Design and immunogenicity of a trihead-I3-A7 nanoparticle immunogen

To evaluate glycosylated I3-A7cp (I3-A7cp-N62) as a nanoparticle vaccine scaffold, we chose to display a domain-based antigen derived from influenza HA. We recently reported the design of RBDs from several HAs stabilized in closed trimeric conformations that we refer to as “triheads” (10, 18). Previously, trihead stabilization required rigid integration into the trimeric component of a two-component protein nanoparticle via a coiled-coil linker comprising a disulfide bond. Displaying the trihead on a different nanoparticle therefore represents a challenging design task.

In parallel to the I3-A7 optimization steps described above, we used RFdiffusion and ProteinMPNN to generate *de novo* linkers between the N terminus of I3-A7cp-N62 and a trihead antigen based on the HA from influenza virus A/Michigan/45/2015 (MI15; (18)). After using RPXdock to obtain an initial dock of the trihead above the I3-A7cp trimer, we provided inputs to RFdiffusion in which the antigen was translated between −6 and +15 Å along the shared three-fold axis of symmetry (**Fig. 5A**). Between five and 80 *de novo* residues were used for linker generation, resulting in primarily alpha-helical structures that were often linked to the N terminus of I3-A7cp by short loops. After designing sequences for the *de novo* linkers using ProteinMPNN, we filtered approximately 2,200 designs based primarily on the accuracy of AF2 prediction of the entire TH-I3-A7 trimer and selected 13 for experimental characterization. Of these, 11 secreted from transfected cells, yielded SEC peaks near the expected elution volume, and bound monoclonal antibodies (mAbs) specific to HA (***SI Appendix*, Fig. S6**). One design, named TH-I3-A7, had 60 residues diffused between the trihead antigen and the underlying nanoparticle, expressed particularly well, and had a nearly symmetric SEC elution peak (**Fig. 5B**). TH-I3-A7 strongly bound the receptor binding site-specific mAb 5J8 (63), and minimally bound the trimer interface-specific mAb FluA-20 (64), indicating display of a closed trihead (**Fig. 5C**). Negative stain EM of TH-I3-A7 revealed fields of monodisperse nanoparticles with visible antigen, and 2D class averages indicated that the *de novo* linker was relatively rigid compared to previously described nanoparticle immunogens that used flexible glycine-serine linkers (**Fig. 5D**; (8)).

**Fig. 5.**
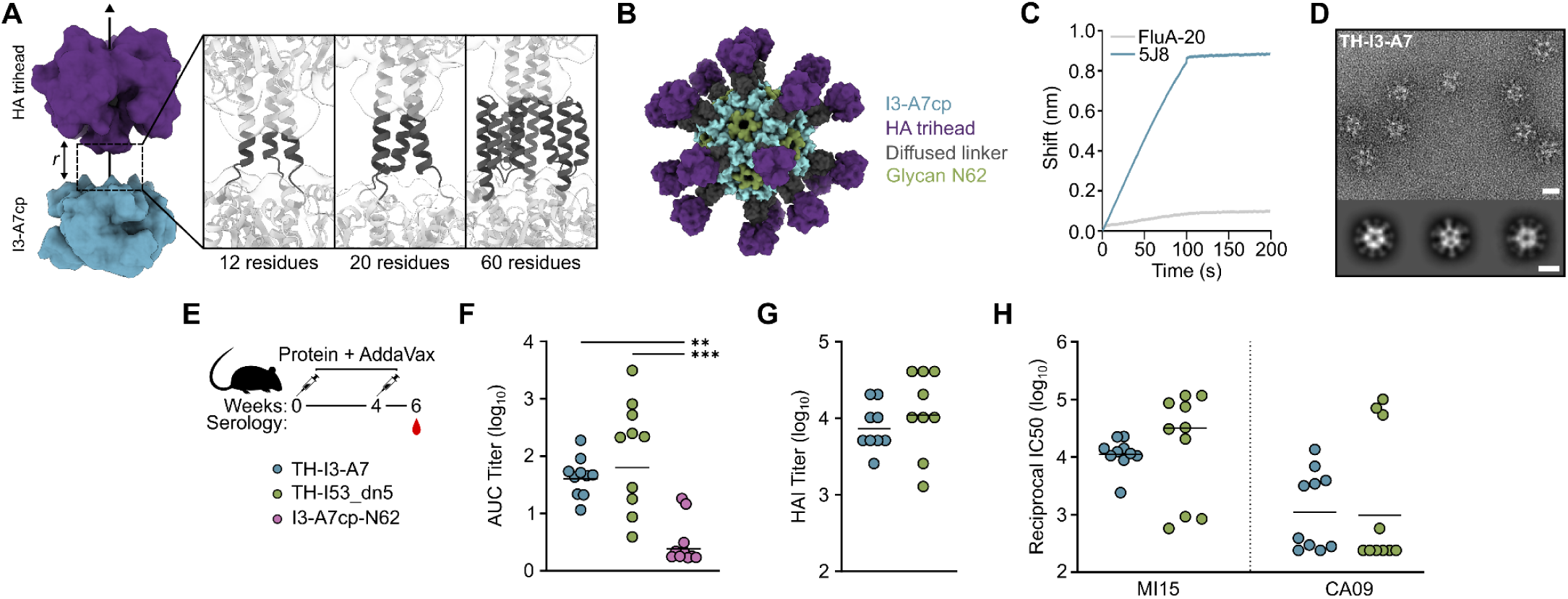
Design and immunogenicity of a trihead-I3-A7 nanoparticle immunogen. *(A) Left,* A schematic highlighting docking of the trihead antigen along the three-fold symmetry axis of the I3-A7cp trimer. Docks with various distances r were provided as inputs to RFdiffusion for linker design. *Right,* Examples of de novo linkers connecting the trihead antigen to the nanoparticle. *(B)* A model of the final nanoparticle vaccine candidate, TH-I3-A7. The I3-A7cp component is shown in blue, the trihead antigen in purple, the oligomannose-type glycan at N62 in green, and the diffused linker in dark gray. *(C)* BLI of the nanoparticle showing strong binding of 5J8 to the receptor binding site and minimal binding of the trimer interface antibody FluA-20. *(D)* A representative electron micrograph and 2D class averages of TH-I3-A7. Scale bar = 40 nm for the micrograph and 20 nm for the 2D class averages. *(E)* TH-I3-A7 immunogenicity study design and groups. *(F)* ELISA binding titers (n=10), *(G)* HAI titers (n=9) and *(H)* microneutralization (n=10) titers in immune sera at week 6. All statistical significance between groups is shown. Significance was determined using the Kruskal-Wallis test corrected with Dunn’s multiple comparisons test (F) or the Mann-Whitney test (G,H); **p < 0.01; ***p < 0.001.

We then evaluated the immunogenicity of TH-I3-A7 in mice, benchmarking against our previously described two-component nanoparticle immunogen displaying the same MI15 trihead (TH-I53_dn5; (18)). We included I3-A7cp-N62 lacking fused antigen as a negative control immunogen. All three immunogens formed monodisperse nanoparticles, and both of the trihead-bearing nanoparticles bound 5J8 but not FluA-20 as shown previously (***SI Appendix*, Fig. S7**). Groups of 10 BALB/c mice were immunized with AddaVax-adjuvanted immunogens comprising 1 μg of trihead or a corresponding number of bare I3-A7cp-N62 nanoparticles at weeks 0 and 4, and sera were collected at week 6 (**Fig. 5F**). The geometric mean titers of HA-specific IgG measured by ELISA were similar between the TH-I3-A7 and TH-I53_dn5 groups, although more variability was observed among the mice that received TH-I53_dn5 (**Fig. 5F**). I3-A7cp-N62 elicited essentially no HA-specific antibodies as expected. We also measured vaccine-matched hemagglutination inhibition (HAI) and neutralization to evaluate the functionality of antibodies elicited by each immunogen. Both assays again showed comparable activity induced by TH-I3-A7 and TH-I53_dn5, with more variability in the TH-I53_dn5 group (**Fig. 5G,H**). Finally, we measured neutralization of the vaccine-mismatched H1N1 strain A/California/07/2009 and similarly found minimal differences between the two groups (**Fig. 5H**). Together, these data establish that the methodologies and design approaches we used were successful in generating an immunogenic nanoparticle vaccine starting from the predicted structure of a thermostable trimeric protein building block.

## Discussion

Here we developed a new method that allowed us to accurately design self-assembling protein nanoparticles using only the amino acid sequences of homologs of oligomeric proteins. The present approach goes beyond previous methods for designing self-assembling proteins by eliminating their requirement for experimentally determined structures of candidate protein building blocks (27–29). This advance was made possible by ML-based tools for structure prediction that can rival the accuracy of experimental structure determination (39, 42, 43). When used as part of the approach we describe here, these tools broaden the number of protein building blocks that can be used to design novel self-assembling proteins, improving the likelihood of obtaining protein assemblies with the desired structural or functional features. Remarkably, the success rate and accuracy of our method (6/88 designs [7%] and 60-subunit alpha-carbon RMSDs of 1.6–2.2 Å, respectively) are comparable to previous reports that started from crystal structures of oligomeric building blocks (27, 29, 47), highlighting the accuracy of both AF2-based structure prediction and the “dock-and-design” methodology. One limitation of the present study is that we only targeted a single symmetric architecture: icosahedral assemblies constructed from trimeric building blocks. However, given that the dock-and-design strategy has successfully generalized to a wide variety of both finite and unbounded architectures (30–32), we expect the present approach to be at least as widely applicable, if not more so due to the increased variety of candidate building blocks.

Several properties are required of a protein assembly in order for it to be a successful one-component nanoparticle vaccine scaffold. This fact was highlighted by our observation that I3-A7 was the only one out of six designed nanoparticles that secreted from mammalian cells, as well as the extensive re-engineering required to transform it into a functional vaccine candidate. We recently showed that the presence of cryptic transmembrane domains in designed protein assemblies can prevent secretion, which prompted the development of a computational protocol to eliminate them through sequence redesign (31). However, none of the six nanoparticles designed in the present study comprised cryptic transmembrane domains, indicating that other unidentified features must account for the failure of five of the six to secrete. As additional determinants of secretion are identified, they can be incorporated into computational design approaches to ensure optimization of this critical property. We then had to substantially alter the structure of I3-A7 to enable display of the trihead antigen by relocating its N terminus and designing a *de novo* linker between the antigen and nanoparticle scaffold. These alterations were necessary because we did not preselect building blocks with particular geometric features. Future efforts can now do so, a strategy that will be facilitated by the wider variety of building blocks now accessible for nanoparticle design. Nevertheless, we were able to successfully re-engineer I3-A7 to display the trihead, a challenging design task that was made possible because of new tools for protein design including RFdiffusion, ProteinMPNN, and AF2. As these tools continue to improve, designing new nanoparticles and manipulating existing ones should become more straightforward. Finally, we engineered a genetically encoded oligomannose-type glycan into the I3-A7 subunit. A limitation of our approach is that we did not explicitly model the glycan during design, as available tools for glycan modeling were few and unwieldy. Versions of both AlphaFold and RoseTTAFold are now available that explicitly model non-protein atoms including glycans (65, 66), and we anticipate that these tools will enable more sophisticated glycan modeling and design. Even without these newer tools, we succeeded in encoding an oligomannose-type glycan in the amino acid sequence of I3-A7, and showed that its composition depended on nanoparticle assembly. To our knowledge, this is the first evidence of the assembly of a secreted protein nanoparticle immunogen during the early stages of the secretory pathway.

Immunogenicity studies in mice showed that TH-I3-A7 is immunogenic and induces robust levels of receptor-blocking and neutralizing antibodies. Notably, TH-I3-A7 is as immunogenic as the same antigen displayed on I53_dn5, the two-component protein nanoparticle on which the trihead was originally designed (10). Two-component nanoparticles like I53_dn5 and homomeric nanoparticles like I3-A7 each have advantages and disadvantages as vaccine platforms. The ability to assemble two-component nanoparticles *in vitro* yields superior control over the quality and homogeneity of the displayed antigen (11) and simplifies the generation of mosaic nanoparticle immunogens that co-display multiple different antigens on the same nanoparticle (23, 67–69). Furthermore, *in vitro* assembly has been de-risked as a scalable manufacturing approach by several two-component nanoparticle immunogens that have advanced to clinical trials or licensure, including multivalent influenza vaccine candidates based on I53_dn5 (NCT06481579, NCT05968989, and NCT04896086; (4, 8, 23, 24)). However, two-component nanoparticles have not yet been adapted for use in genetic immunization approaches based on DNA, mRNA, or viral vectors. These can be faster to manufacture than protein-based vaccines, which is particularly relevant in indications like influenza where frequent strain updates are required due to antigenic drift and shift (70). Several homomeric protein nanoparticle immunogens have been delivered as either DNA or mRNA vaccines, mostly based on a small number of naturally occurring self-assembling proteins such as ferritin and lumazine synthase (71–75). Computationally designing novel homomeric nanoparticles promises to enhance the performance of genetically delivered nanoparticle vaccines by enabling precise control over their size, shape, symmetry, valency, and other features. By broadening the set of oligomeric protein building blocks available for design, our method will facilitate efforts directed at designing self-assembling protein nanomaterials with structures and activities tailored for this and other applications in medicine.

### Data, Materials, and Software Availability

Data deposition, atomic coordinates, and structure factors have been deposited in the Protein Data Bank, http://www.rcsb.org (PDB ID 9CLZ, 9CM1, and 9CM0). Cryo-EM maps were deposited in the Electron Microscopy Data Bank (EMD-45734, EMD-45736, and EMD-45735). These data will be released prior to publication.

## Methods

### Identifying and filtering sequences from thermophiles

To identify initial structures on which to base homology, the PDB was searched for homotrimers with C3 symmetry at a sufficient resolution (<2.4 Å). Additional filters ensured that selected proteins: (1) maintain >50% helical structure for easier implementation with current nanoparticle docking algorithms; (2) originate in Archaea or Bacteria so that they might have thermophilic homologs; and, (3) do not include membrane proteins to reduce the risk of insoluble protein expression *in vitro*. The resulting list of PDB seeds was used to search the UniProt database for all sequences with >50% identity to each seed. For those sequences, the parent organisms were considered thermophiles if “therm” was in the name or if they had reported optimal growth temperatures above 55°C on the BacDive database. Remaining hits were filtered to <70% identity so sequences were not too similar. Sequences with published high-resolution structures were also removed.

### Computational design of protein nanoparticles

Thermophilic C3 sequences were predicted using AlphaFold2-ptm and predictions were filtered to keep only those with a predicted local distance difference test (pLDDT) > 90 and a predicted aligned error (PAE) < 5. Asymmetric units for each homotrimer were extracted and used to establish an initial nanoparticle docking conformation using RPXDock. Further sampling was introduced with Rosetta SymDofMover by translating trimers along the radial axis and rotating them about their symmetry axis. The interface residues were defined as those within 8 Å of each other at contact sites between components. Designs were eliminated if there were less than 12 interface residues. For all other designs, ProteinMPNN was implemented with a sampling temperature of 0.2 to engineer the interface residues for optimal self-assembly. After ten iterations of ProteinMPNN for each interface, the sequences were threaded onto the AF2 predictions with RosettaScripts and Rosetta metrics were calculated for filtering purposes. The following filters eliminated most designs: ddG < −10, Solvent Accessible Surface Area (SASA) > 300, Shape Complementarity > 0.5, and Degreaser Metric > 2.6. Prior to ordering constructs, the predicted sequences (which were derived from the originating PDB seeds and comprised only the regions modeled in the structures) were used as inputs in BLAST (76) to retrieve full-length genomic sequences, and these were used as the basis for construct generation.

### Bacterial protein expression and purification

To screen initial designs, eBlocks gene fragments (IDT) were cloned into a pET29b+ based vector backbone via Golden Gate Assembly and transformed into chemically competent BL21 *E. coli* (NEB). Small scale production occurred in 96 deep-well plates in autoinduction media containing TBII media (Mpbio) supplemented with 50✕5052, 50✕M salts, 20 mM MgSO_4_, and a trace metal mix. Cultures grew under antibiotic selection at 37°C for 6 hours at 1350 rpm in a Heidolph shaker. Cells were harvested by centrifugation at 4,000 *x g* and chemically lysed with BugBuster protein extraction reagent (Sigma-Aldrich) after resuspension in lysis buffer (50 mM Tris pH 8.0, 250 mM NaCl, 20 mM imidazole), followed by addition of bovine pancreatic DNaseI (Sigma-Aldrich) and protease inhibitors (Thermo Scientific). Clarified lysate supernatants were applied to equilibrated Ni-NTA resin (QIAGEN) columns. Washes were performed with 10 column volumes of lysis buffer, then eluted with 4 column volumes of the same buffer containing 500 mM imidazole. Eluant was analyzed for protein expression via SDS-PAGE and HPLC.

Large-scale nanoparticle expression (50-100 mL) was performed as described above, except using TBII media with IPTG induction at OD_600_=0.6-0.8 and 4 hours of expression at 37°C and 250 rpm. Concentrated or unconcentrated elution fractions were further purified using a Superose 6 Increase 10/300 GL (Cytiva) on an ÄKTA Pure (Cytiva) into 25 mM Tris pH 8.0, 150 mM NaCl. Instrument control and elution profiles analysis were performed with UNICORN 7 software (Cytiva).

### Mammalian protein expression and purification

For purification of plasmid DNA for Expi293F transfection, bacteria were cultured and plasmids were harvested according to the QIAGEN Plasmid Plus Maxi Kit protocol (QIAGEN). For mammalian expression and purification of secreted nanoparticles, Expi293F cells at 3 million cells/mL were transfected with the following per mL of cell culture: 70 uL Opti-MEM (Thermo Scientific), 1 µg plasmid DNA and 3 µg PEI-MAX. Cells were harvested 3-5 days post-transfection by centrifugation for 5 minutes at 4,000 *x g*, incubation with polyDADMAC (Sigma-Aldrich) for 10 minutes, and a final centrifugation for 5 minutes at 4,000 *x g*. Sterile-filtered supernatant was adjusted to 50 mM Tris pH 8.0 and 500 mM NaCl, then batch bound to Ni Sepharose Excel (Cytiva) with agitation at 25°C for 1 hour or 4°C for 15 hours. Pelleted resin was washed with 20 column volumes of wash buffer (50 mM Tris pH 8.0, 500 mM NaCl, 30 mM imidazole), then eluted with 2 column volumes of the same buffer containing 300 mM imidazole. Concentrated elution fractions were purified by size-exclusion chromatography as described above, with the addition of 100-200 mM L-Arginine as an excipient for I3-A7cp and constructs fused to antigen. For TH-I3-A7, the mutation T38E (I3-A7cp numbering) was made to improve secretion yields after using DeepTMHMM prediction (77) to identify potential cryptic transmembrane regions.

### DLS and thermal melts

DLS was performed using an UNcle (Unchained Labs) to determine the hydrodynamic radius of each nanoparticle and the polydispersity of the sample. SEC-purified samples at 0.1 mg/mL were loaded into an 8.8 μL quartz capillary cassette (UNi, UNchained Laboratories) and DLS measurements were obtained at 25°C. Buffer components and protein compositions were accounted for in the UNcle software. To determine the aggregation temperature (*T_agg_*) of the nanoparticles, DLS measurements were collected over a temperature range from 25-84°C or 25-95°C, with a ramp of approximately 2°C/min. *T_agg_* was defined as the temperature at which the absolute value of the first derivative of the Z-averaged diameter was greater than unity for all subsequent consecutive measurements while allowing for single nonconsecutive values below one. First derivatives were calculated after smoothing with four neighboring points using 2nd order polynomial settings in GraphPad Prism. Two replicates were performed for each nanoparticle.

### Biolayer interferometry (BLI)

BLI was performed using an Octet Red 96 system, at 25°C and 1000 rpm. Antibodies were diluted to 0.02 mg/mL (5J8) and 0.07 mg/mL (FluA-20) in kinetics buffer (1✕Phosphate-buffered saline (PBS), 0.5% serum bovine albumin and 0.01% Tween). ProA Octet Biosensors (Sartorius) were used to load antibodies for the assay which was run as follows: 60 seconds buffer, 100 seconds loading, 30 seconds buffer, 100 seconds association, 100 seconds dissociation. Biosensors were regenerated using 10 mM Glycine pH 2.5 for 30 seconds, followed by 60 seconds in buffer.

### Negative stain electron microscopy (nsEM)

300-mesh carbon coated grids (Electron Microscopy Sciences) were glow discharged for 30 seconds prior to application of protein samples (3 µL, 0.1 mg/mL) and blotting three times with 2% (w/v) uranyl formate. Data were collected using a 120 kV Talos L120C transmission electron microscope (Thermo Scientific) with a BM-Ceta camera. CryoSPARC was used for CTF correction, particle picking and extraction, and 2D classification.

### Cryo-EM sample preparation, data collection and processing

Cryo-EM samples were prepared on glow-discharged CFLAT 1.2/1.3 holey-carbon grids (Electron Microscopy Sciences). Protein sample at 1-4 mg/ml was applied to each grid, incubated for 30 seconds, and then blotted using Whatman filter paper; this process was performed twice, followed by a third application of sample and final blotting and plunge freezing in liquid-ethane using a Vitrobot (FEI). Data were acquired on a Titan Krios operating at 300 kV equipped with a K3 Summit Direct Detector and a Quantum GIF energy filter (Gatan) operating in zero-loss mode with a 20-eV slit width. Movies were collected in counting mode, collecting 50 frames with a total dose of 52 e−/Å−2, with a defocus range of −0.5 to −2 µm. Automated data collection was carried out using Leginon (78) at a nominal magnification of 105,000× with a pixel size of 0.843 Å. All data processing was carried out using cryoSPARC (79). Movie frame alignment and dose-weighting summation were performed using Patch Motion, and CTF estimation was done using patch CTF. Particles were first picked using Blob picker and these were used to generate initial 2D averages. A few of the best 2D averages representative of different particle views were then selected as templates for template picking. From these template picks, two rounds of 2D classification and selection were done to isolate well-aligned, single particles. Initial models were generated by ab-initio reconstruction with C1 symmetry. Homogeneous 3D refinements were then carried out with icosahedral symmetry, with per-particle defocus refinement, and beamtilt and anisotropic magnification by image shift group enabled.

### Model building and refinement

AlphaFold2 models were docked into cryo-EM maps as rigid bodies using Chimera (80). An initial refinement of a nanoparticle asymmetric unit (asu) into the cryo-EM density was performed with ISOLDE (81). A full icosahedron was then generated from the ISOLDE-refined asu, and then this entire icosahedral nanoparticle was relaxed into the cryo-EM density again in ISOLDE. A final real-space refinement with grid searching, Ramachandran and rotamer restraints all turned off, and ‘use starting model as reference’ turned on, was then performed in Phenix (82). There were flexible loops comprised of residues 17-23 and 106-113 in I3-A6, residues 123-129 in I3-A7, and residues 137-143 and 170-176 in I3-D12 that were not resolved in the cryo-EM density so these loops were not modeled.

### Sample preparation and analysis by LC-MS

Glycan analysis was performed as described previously (83–85). Briefly, aliquots of each sample were denatured for 1h in 50 mM Tris/HCl, pH 8.0 containing 6 M of urea for 1 hour. The polypeptides were buffer-exchanged into 50 mM Tris/HCl, pH 8.0 using Vivaspin columns (3 kDa) and digested separately overnight using trypsin/Lys-C (1 µg), chymotrypsin (1 µg) or alpha lytic protease (1 µg) (Mass Spectrometry Grade, Promega). The next day, the peptides were dried and extracted using Oasis HLB 96 well plate (Waters). The peptides were analyzed by nanoLC-ESI MS with an Ultimate 3000 HPLC (Thermo Fisher Scientific) system coupled to an Orbitrap Eclipse mass spectrometer (Thermo Fisher Scientific) using stepped higher energy collision-induced dissociation (HCD) fragmentation (15%, 25%, 45%). Peptides were separated using an EasySpray PepMap RSLC C18 column (75 µm × 15 cm). A trapping column (PepMap 100 C18 3 μm particle size, 75 μm x 2 cm) was used in line with the LC prior to separation with the analytical column. The LC conditions were as follows: 60-minute linear gradient consisting of 5-40% acetonitrile in 0.1% formic acid over 50 minutes followed by 10 minutes of 95% acetonitrile in 0.1% formic acid. The flow rate was set to 300 nL/min. The spray voltage was set to 2.5 kV and the temperature of the heated capillary was set to 55 °C. The ion transfer tube temperature was set to 275 °C. Precursor and fragment detection were performed using an Orbitrap at a resolution MS1 = 120,000, MS2 = 30,000. Glycopeptide fragmentation data were extracted from the raw file using Byos (version 5.5, Protein Metrics). The glycopeptide fragmentation data were evaluated manually for each glycopeptide; the peptide was scored as true-positive when the correct b and y fragment ions were observed along with oxonium ions corresponding to the glycan identified. The MS data was searched using the Protein Metrics N-glycan library. The relative amounts of each glycan at each site as well as the unoccupied proportion were determined by comparing the extracted chromatographic areas for different glycotypes with an identical peptide sequence. All charge states for a single glycopeptide were summed. The precursor mass tolerance was set at 4 ppm and 10 ppm for fragments. A 1% false discovery rate (FDR) was applied. Glycans were categorized according to the composition detected. HexNAc(2)Hex(9−4) was classified as M9 to M4. Any of these compositions containing fucose were classified as fucosylated mannose (FM). HexNAc(3)Hex(5−6)X was classified as Hybrid with HexNAc(3)Fuc(1)X classified as Fhybrid. Complex-type glycans were classified according to the number of processed antenna and fucosylation. Complex-type glycans were categorized according to the number of N-acetylhexosamine monosaccharides detected that do not fit in the previously defined categories.

### Immunization

For immunogenicity studies, female BALB/cAnNHsd were purchased from Envigo (order code 047) at 7 weeks of age. Mice were housed in a specific-pathogen free facility within the Department of Comparative Medicine at the University of Washington, Seattle, accredited by the Association for Assessment and Accreditation of Laboratory Animal Care (AAALAC). Animal studies were conducted in accordance with the University of Washington’s Institutional Animal Care and Use Committee under protocol 4470-01. For each immunization, low-endotoxin proteins were diluted to 20 μg/mL in buffer and mixed with 1:1 v/v AddaVax adjuvant (InvivoGen vac-adx-10) to obtain a final dose of 1 μg of immunogen per animal, per injection. At 8 weeks of age, 10 mice per group were injected subcutaneously in the inguinal region with 100 μL of immunogen at weeks 0 and 4. Animals were bled using the submental route at week 6. Whole blood was collected in serum separator tubes (BD #365967) and rested for 30 min at room temperature for coagulation. Tubes were then centrifuged for 10 min at 2,000 x *g* and serum was collected and stored at −80°C until use.

### Enzyme-linked immunosorbent assay (ELISA)

HA-foldon trimers were added to 96-well Nunc MaxiSorp plates (Thermo Scientific) at 2.0 μg/mL with 100 mL per well and incubated for 18 h. Blocking buffer (TBST: 25 mM Tris pH 8.0, 150 mM NaCl, 0.05% (v/v) Tween 20, and 5% Nonfat milk) was then added at 200 mL per well and incubated for 1 h. Plates were then washed three times with TBST using a robotic plate washer (Biotek). Three-fold serial dilutions of serum starting at 1:100 were made in blocking buffer and added to plates at 100 μL per well, and incubated for 1 h. Plates were washed thrice again before addition of 100 μL per well of anti-mouse IgG HRP-linked secondary antibody (CellSignaling Technology) diluted 1:2,000 in blocking buffer and incubated for 30 min. All incubations were carried out with shaking at room temperature. Plates were washed a final time, and then 100 μL per well of TMB (3,30,50,5-tetramethylbenzidine, SeraCare) was added for 2 min, followed by quenching with 100 μL per well of 1 N HCl. Reading at 450 nm absorbance was done on an Epoch plate reader (BioTek).

### Hemagglutination inhibition (HAI)

Serum was inactivated using receptor-destroying enzyme (RDE) II (Seiken) in PBS at a 3:1 ratio of RDE to serum for 18 h at 37°C, followed by 40 min at 56°C. Inactivated serum was serially diluted 2.5-fold in PBS in V-bottom plates at 25 μL per well. 25 μL HA-ferritin nanoparticles at 4 hemagglutinating units were then added to all wells and incubated at room temperature for 30 min. Lastly, 50 μL of 10-fold diluted turkey red blood cells (Lampire) in PBS was added to each well. Hemagglutination was left to proceed for at least 1 h before recording HAI titer.

### Reporter-based microneutralization assay

Microneutralization assays were performed as described previously (86). Generation of the replication-restricted reporter (R3ΔPB1) virus H1N1 A/Michigan/45/2015 and A/California/07/2009 is described elsewhere (87). Briefly, to produce the R3 viruses, the viral genomic RNA encoding functional PB1 was replaced with a gene encoding the fluorescent protein (TdKatushka2), and the R3 viruses were rescued by reverse genetics and propagated in the complementary cell line which expresses PB1 constitutively. Each R3 virus stock was titrated by determining the fluorescent units per mL (FU/mL) prior to use in the experiments. For virus titration, serial dilutions of virus stock in OptiMEM were mixed with pre-washed MDCK-SIAT1-PB1 cells (8 x 10^5^ cells/ml) and incubated in a 384-well plate in quadruplicate (25 µL/well). Plates were incubated for 18-26 h at 37°C with 5% CO2 humidified atmosphere. After incubation, fluorescent cells were imaged and counted by using a Celigo Image Cytometer (Nexcelom) with a customized red filter for detecting TdKatushka2 fluorescence.

For the microneutralization assay, serial dilutions of RDE (Receptor Destroying Enzyme) treated serum were prepared in OptiMEM and mixed with an equal volume of R3 virus (∼8 x 10^4^ FU/mL) in OptiMEM. After incubation at 37°C and 5% CO2 humidified atmosphere for 1 h, pre-washed MDCK-SIAT1-PB1 cells (8 x 10^5^ cells/well) were added to the serum-virus mixtures and transferred to 384-well plates in quadruplicate (25 µL/well). Plates were incubated and counted as described above. Target virus control range for this assay is 500 to 2,000 FU per well, and cell-only control is acceptable up to 30 FU per well. The percent neutralization was calculated for each well by constraining the virus control (virus plus cells) as 0% neutralization and the cell-only control (no virus) as 100% neutralization. A 7-point neutralization curve was plotted against serum dilution for each sample, and a four-parameter nonlinear fit was generated using Prism (GraphPad) to calculate the 50% (IC50) inhibitory concentrations.

## Acknowledgements

The authors thank Andrew Borst, Kenneth Carr, and Connor Weidle for assistance with electron microscopy; Suna Cheng, Craig Dobbins, Alexis Hand, and Adri Tran-Pearson for assistance with tissue culture; Lauren Carter and Kandise VanWormer for laboratory and administrative support; and Luki Goldschmidt and Patrick Vecchiato for building and maintaining the computing infrastructure at the Institute for Protein Design. This work was supported by a generous gift from Open Philanthropy; the Audacious Project at the Institute for Protein Design; the Bill & Melinda Gates Foundation (INV-010680 and INV-043758); the IAVI Neutralizing Antibody Center through the Collaboration for AIDS Vaccine Discovery grant OPP1196345/INV-008813 funded by the Bill & Melinda Gates Foundation; the National Institute of Allergy and Infectious Diseases (P01AI167966); the intramural research program of the Vaccine Research Center, National Institute of Allergy and Infectious Diseases, National Institutes of Health; and the National Science Foundation Graduate Research Fellowships Program Award (DGE-2140004 to C.M.H.).

