## SupplementaryData for "From sequence to scaffold: computational design of protein nanoparticle vaccines from AlphaFold2-predicted building blocks"

**This PDF file includes:**

SI Figures S1 to S8
SI Tables S1 to S3
SI References

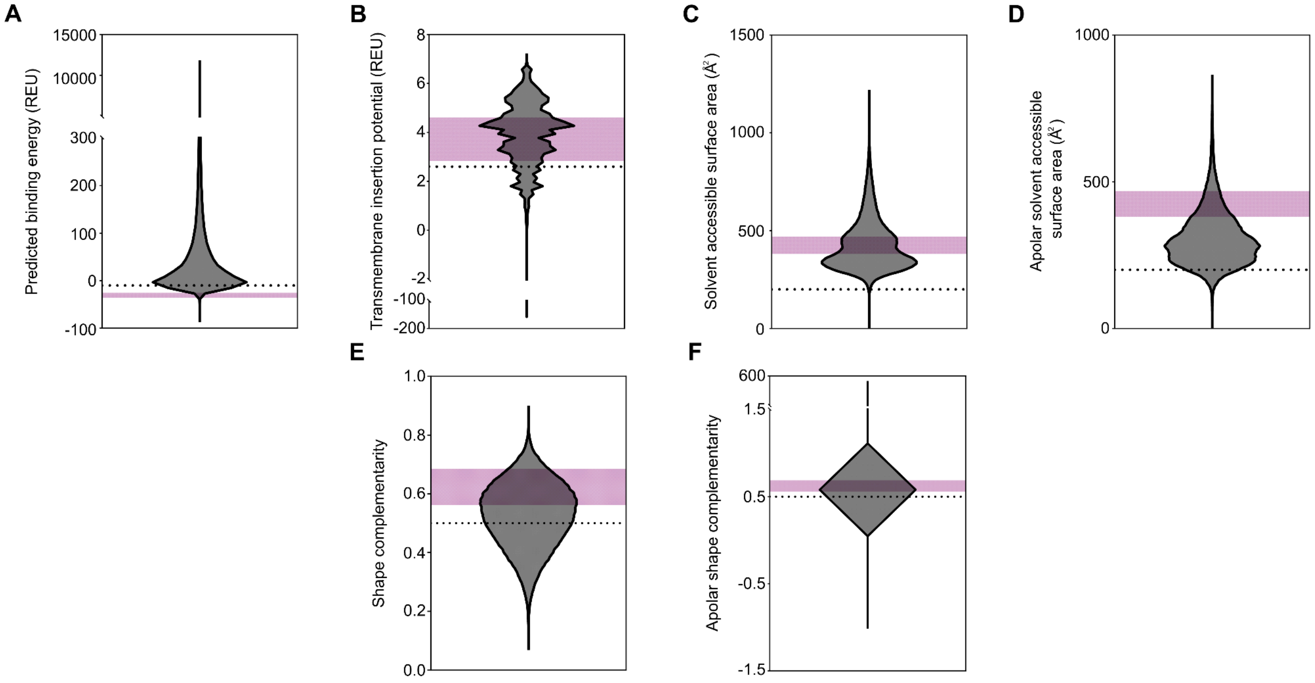


**Fig. S1. Summary of Rosetta scoring metrics used to filter and select final nanoparticle designs.**
The distribution of data points for the metrics used to filter all designs: *(A)* predicted binding energy (ddG), *(B)* transmembrane insertion potential, *(C)* buried solvent accessible surface area, *(D)* buried apolar solvent accessible surface area, *(E)* shape complementarity, and *(F)* apolar shape complementarity. A small number of values less than zero for buried solvent accessible surface area were attributed to Rosetta calculation errors and are not shown. The dotted lines represent cutoffs used and the magenta regions identify the ranges within which the successfully assembling nanoparticles were selected from.

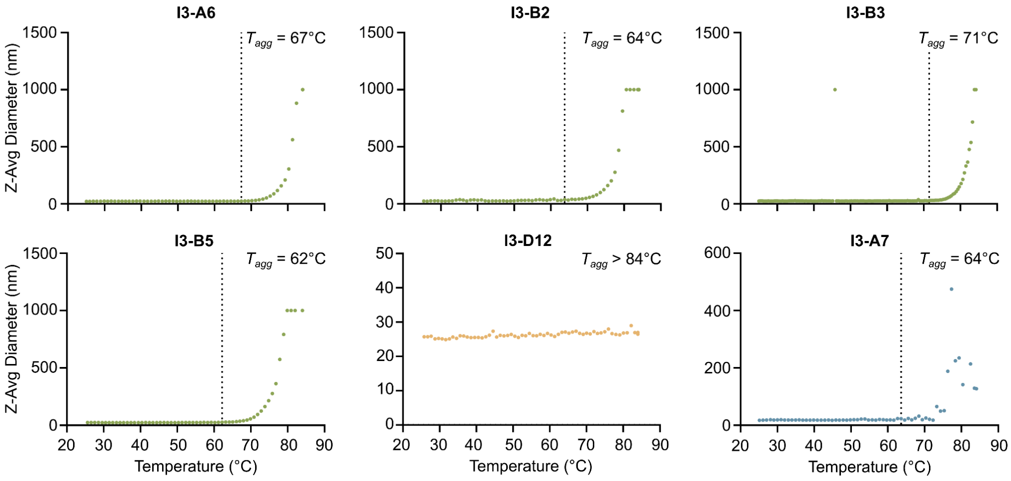


**Fig. S2. DLS melting curves used to determine aggregation temperature of nanoparticles.** The Z-average diameter of each nanoparticle as determined by DLS with a thermal ramp.

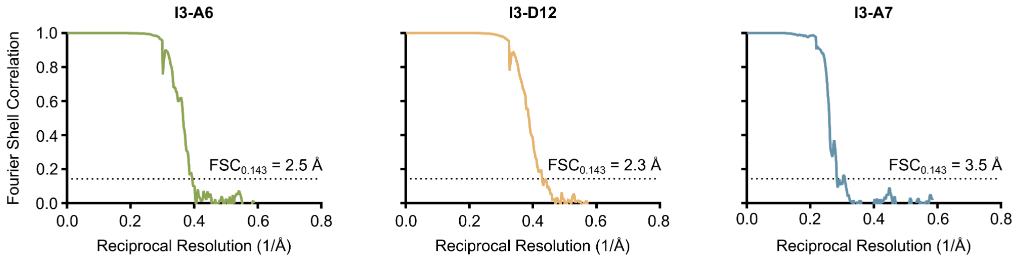


**Fig. S3. Cryo-EM gold-standard Fourier shell correlation (FSC) curves.**
Gold-standard FSC curves for cryo-EM nanoparticle reconstructions. The 0.143 cutoff is indicated by the dashed line.

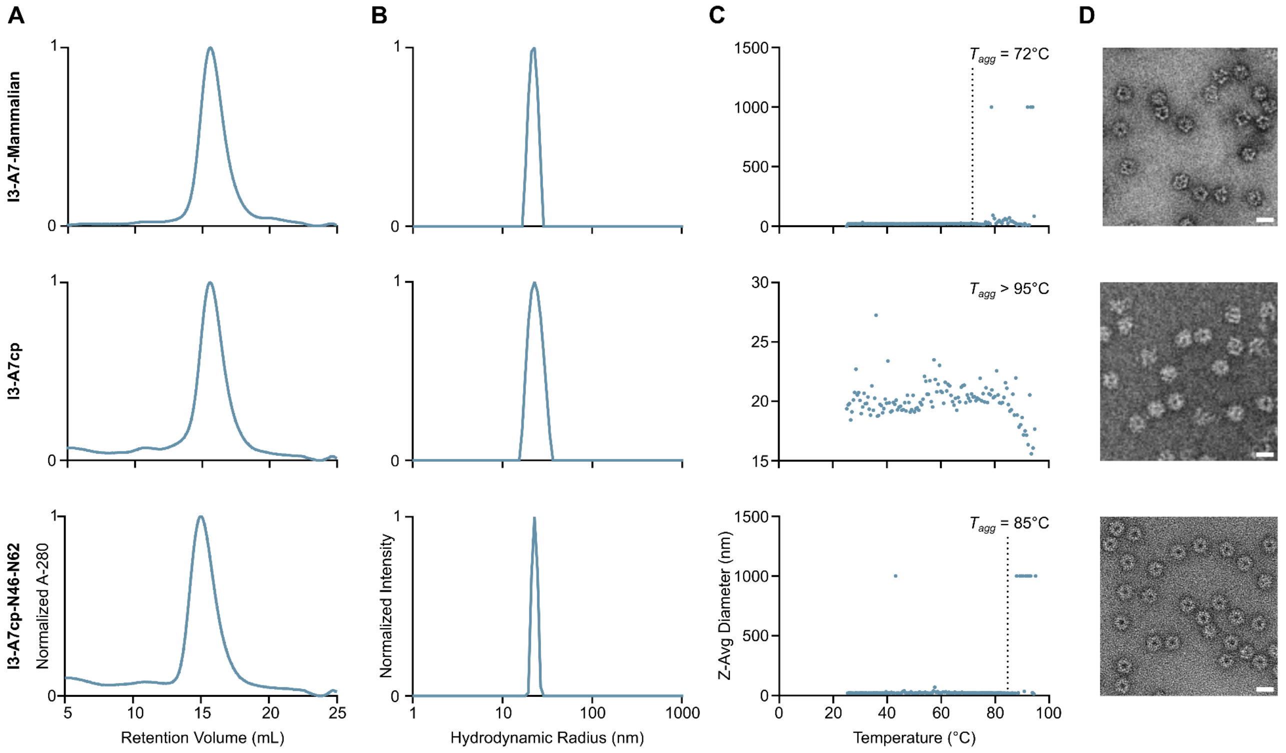


**Fig. S4. Characterization of each variant of nanoparticle I3-A7.**
*(A)* SEC traces, *(B)* DLS, *(C)* thermal melting curves, and *(D)* negative stain electron micrographs for I3-A7 secreted from mammalian cells, the circular permuted version (I3-A7cp), and the permutation with the two N-linked glycans at N46 and N62. Micrographs for I3-A7cp and I3-A7cp-N46-N62 are differently cropped images of those shown in Figure 4. Scale bar = 21 nm.

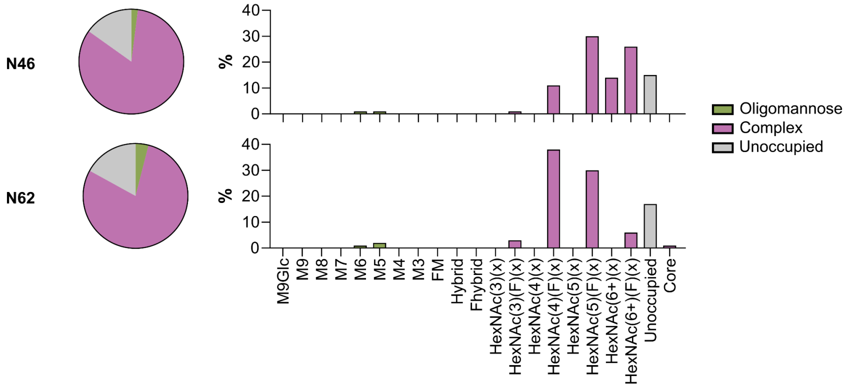


**Fig. S5. Characterization of glycans N46 and N62 in the I3-A7cp-N46-N62-NonAssembling trimer.**
The glycan composition of peptides containing the N46 (top) and N62 (bottom) glycosylation sites of the I3-A7cp-N46-N62-NonAssembling trimer determined by LC-MS.

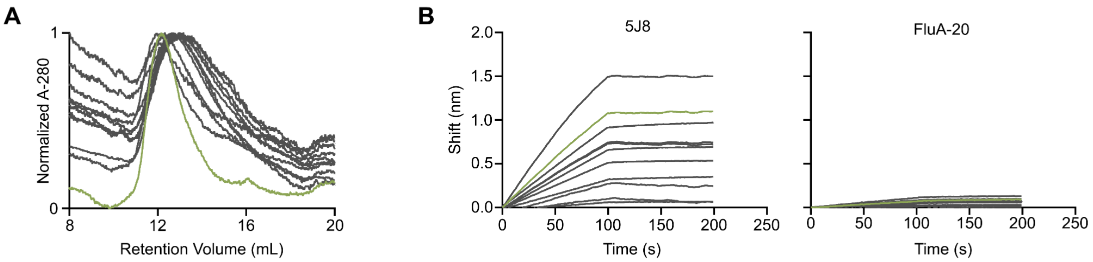


**Fig. S6. Experimental characterization of trihead-bearing I3-A7cp nanoparticles with eleven diffused linkers by SEC and BLI.**
*(A)* SEC traces of the eleven potential vaccine candidates. *(B)* BLI of nanoparticle vaccine candidates binding to 5J8 and FluA-20. The green line in all plots signifies the final candidate tested *in vivo* (TH-I3-A7).

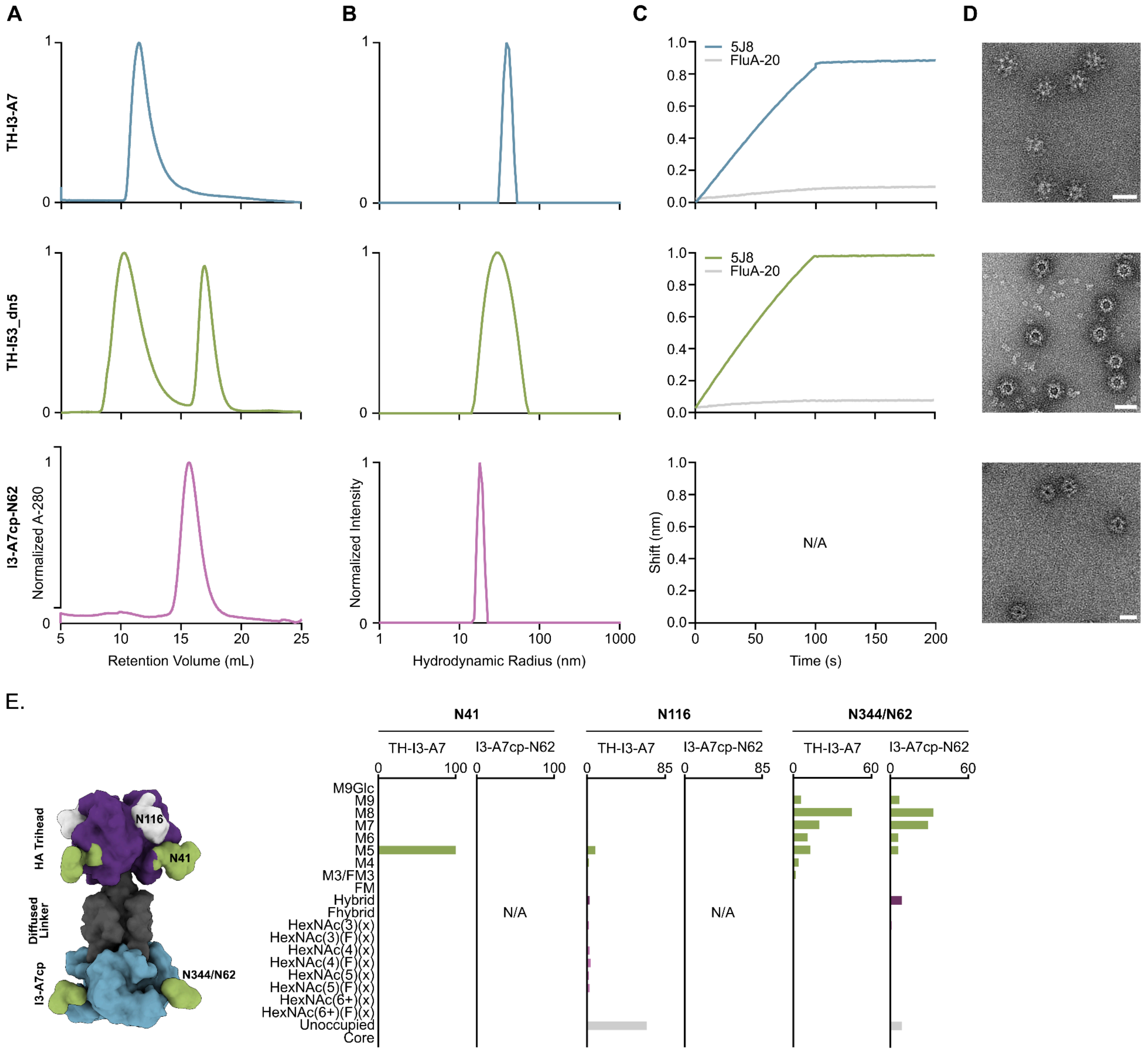


**Fig. S7. Biochemical, biophysical, and antigenic characterization of nanoparticle immunogens.**
*(A)* SEC traces, *(B)* DLS, *(C)* BLI, and *(D)* negative stain EM of the three immunogens used in the mouse immunogenicity study shown in Figure 5. Scale bar = 40 nm, 42 nm, and 21 nm for TH-I3-A7, TH-I53_dn5, and I3-A7cp-N62, respectively. *(E)* Characterization of the three primary glycosylation sites on TH-I3-A7. N41 and N116 are in the trihead antigen, and N344/N62 is in I3-A7cp. BLI and negative stain EM for TH-I3-A7 are shown here in addition to Figure 5 for comparison.

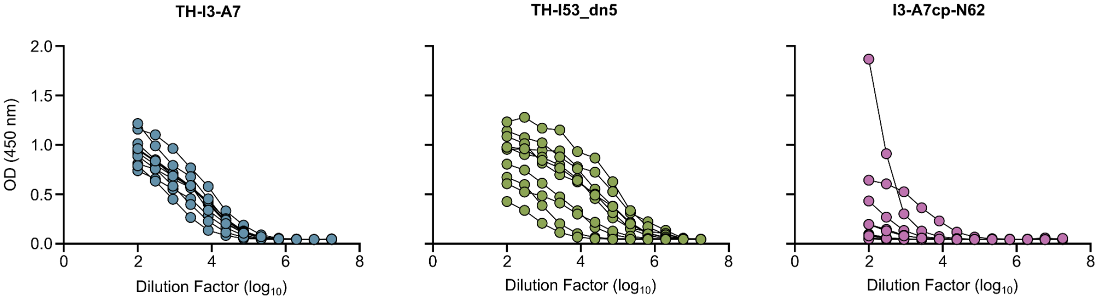


**Fig. S8. ELISA curves for mouse serum antibody binding to a full-length MI15 ectodomain trimer.**
Week six serum ELISA curves against a vaccine-matched full-length HA ectodomain trimer (A/Michigan/45/2015).

**Table S1. Interface Mutations of Designed Nanoparticles**

| Nanoparticle | PDB Seed | Homolog Source | Homolog Sequence | Nanoparticle Sequence |
|---|---|---|---|---|
| I3-A6 | 6HS8 | *Clostridium sp.* CAG:307_30_263; Genbank Accession OKZ90856 | MKILVINGPNINFLGIREKG<br>IYGNDNYDTLVDMINKKAE<br>ELNVKVEVFQSNHEGAIID<br>KLQEAYYNDVDGIVINPGA<br>FTHYSYAVRDALASIAAIP<br>KIEVHISNVHTREEFRHTS<br>VTVPVCQGQVVGLGLRG<br>YLYAMEALTDMTKSN | MKILVINGPNINFLGIREKG<br>IYG**PL**NYD**D**LV**E**MI**KGT**A**K**<br>**G**L**K**VKVEVFQSNHEGAIID<br>KLQEAYYNDVDGIVINPGA<br>FTHYSYAVRDALASIAAIP<br>KIEVHISNVHTREEFRHTS<br>VTVPVCQG**E**VVGLGL**G**G<br>YL**A**AM**GM**L**VE**MTKSN |
| I3-A7 | 6HF7 | *Thermoplasmata archaeon HGW-Thermoplasmata -1*; Genbank Accession PKK86397 | MGAIVVTGIPGVGKTTVM<br>QKAAEGMNIEFVTYGTEM<br>FEVAKSMGLVKDRDEMR<br>RLSPDVQKEVQKKAAENI<br>AAKGNVILDTHCTIKTPKG<br>YLPGLPRWVLEKLRPSVIL<br>LVEADPKEIYGRRLKDET<br>RNRDPDSEEEIAEHQMM<br>NRAAAMAYASLSGATVKIV<br>FNHDNRLDDAVRDAAPVL<br>GN | MGAIVVTGIPGVGKTTVM<br>QKAAEG**SPLPR**V**PLEGV**<br>M**YG**VAK**R**MGLVKD**I**DEMR<br>RLSPDVQKEVQKKAAE**R**I<br>AA**L**G**D**VILDTHCTIKTPKG<br>YLPGLPRWVLEKLRPSVIL<br>LVEADPKEIYGRRLKDET<br>RNRDPDSEEEIAEHQMM<br>NRAAAMAYASLSGATVKIV<br>FNHDNRLDDAVRDAAPVL<br>GN |
| I3-B2 | 6HS8 | *Clostridium sp.* CAG:307_30_263; Genbank Accession OKZ90856 | MKILVINGPNINFLGIREKG<br>IYGNDNYDTLVDMINKKAE<br>ELNVKVEVFQSNHEGAIID<br>KLQEAYYNDVDGIVINPGA<br>FTHYSYAVRDALASIAAIP<br>KIEVHISNVHTREEFRHTS<br>VTVPVCQGQVVGLGLRG<br>YLYAMEALTDMTKSN | MKILVINGPNINFLGIREKG<br>IYG**KY**NYD**F**LVD**L**IN**ET**A**K**<br>**L**LNVKVEVFQSNHEGAIID<br>KLQEAYYNDVDGIVINPGA<br>FTHYSYAVRDALASIAAIP<br>KIEVHISNVHTREEFRHTS<br>VTVPVCQG**EID**G**A**GL**G**GY<br>**VS**AM**LT**L**VE**MTKSN |
| I3-B3 | 6HS8 | *Clostridium sp.* CAG:307_30_263; Genbank Accession OKZ90856 | MKILVINGPNINFLGIREKG<br>IYGNDNYDTLVDMINKKAE<br>ELNVKVEVFQSNHEGAIID<br>KLQEAYYNDVDGIVINPGA<br>FTHYSYAVRDALASIAAIP<br>KIEVHISNVHTREEFRHTS<br>VTVPVCQGQVVGLGLRG<br>YLYAMEALTDMTKSN | MKILVINGPNINFLGIREKG<br>IYG**PF**NYD**F**LV**EL**INK**T**AE<br>**L**LNVKVEVFQSNHEGAIID<br>KLQEAYYNDVDGIVINPGA<br>FTHYSYAVRDALASIAAIP<br>KIEVHISNVHTREEFRHTS<br>VTVPVCQG**E**V**D**G**A**GL**G**G<br>Y**VS**AME**R**L**VS**MTKSN |
| I3-B5 | 6HS8 | *Clostridium sp.* CAG:307_30_263; Genbank Accession OKZ90856 | MKILVINGPNINFLGIREKG<br>IYGNDNYDTLVDMINKKAE<br>ELNVKVEVFQSNHEGAIID<br>KLQEAYYNDVDGIVINPGA<br>FTHYSYAVRDALASIAAIP<br>KIEVHISNVHTREEFRHTS<br>VTVPVCQGQVVGLGLRG<br>YLYAMEALTDMTKSN | MKILVINGPNINFLGIREKG<br>IYG**PF**NYD**D**LVD**L**INK**T**AE<br>**F**LNVKVEVFQSNHEGAIID<br>KLQEAYYNDVDGIVINPGA<br>FTHYSYAVRDALASIAAIP<br>KIEVHISNVHTREEFRHTS<br>VTVPVCQGQV**D**G**A**GL**A**G<br>Y**VQ**AM**FE**LT**S**MTKSN |

| | | | | |
|---|---|---|---|---|
| I3-D12 | 1QWG | *Methanocaldococcus jannaschii*; Genbank Accession HII60133 | MKAFEFLYEDFQRGLTVV<br>LDKGLPPKFVEDYLKVCG<br>DYIDFVKFGWGTSAVIDR<br>DVVKEKINYYKDWGIKVY<br>PGGTLFEYAYSKGKFDEF<br>LNECEKLGFEAVEISDGS<br>SDISLEERKNAIKRAKDNG<br>FMVLTEVGKKMPDKDKQL<br>TIDDRIKLINFDLDAGADY<br>VIIEGRESGKGIGLFDKEG<br>KVKENELDVLAKNVDINK<br>VIFEAPQKSQQVAFILKFG<br>SSVNLANIAFDEVISLETL<br>RRGLRGDTFGKV | MKAFEFLYEDFQRGLTVV<br>LDKGLPPKFVEDYLKVCG<br>DYIDFVKFGWGTSAVIDR<br>DVVKEKINYYKDWGIKVY<br>PGGTLFEYAYSKG**EAA**EF**I**<br>**A**EC**K**KLGFEAVEISDGSS<br>DISL**DD**RK**A**AI**WG**AK**WA**G<br>FMVLTEVGKKMPDKDKQL<br>TIDDRIKLINFDLDAGADY<br>VIIEGRESGKGIGLFDKEG<br>KVKENELDVLAKNVDINK<br>VIFEAPQKSQQVAFILKFG<br>SSVNLANIAFDEVISLETL<br>RRGLRGDTFGKV |

Bolded and underlined letters in the nanoparticle sequences represent the mutations designed for nanoparticle assembly.

**Table S2. Amino Acid Sequences**

| Construct | Sequence |
|---|---|
| I3-A6 | *MTDEPVATEEPAARDEVRRAEALGGQAAEQLA*MKILVINGPNINFLGIREKGIYGP<br>LNYDDLVEMIKGTAKGLKVKVEVFQSNHEGAIIDKLQEAYYNDVDGIVINPGAFTHY<br>SYAVRDALASIAAIPKIEVHISNVHTREEFRHTSVTVPVCQGEVVGLGLGGYLAAM<br>GMLVEMTKSN**GSWGSLEHHHHHH** |
| I3-B2 | *MTDEPVATEEPAARDEVRRAEALGGQAAEQLA*MKILVINGPNINFLGIREKGIYGK<br>YNYDFLVDLINETAKLLNVKVEVFQSNHEGAIIDKLQEAYYNDVDGIVINPGAFTHY<br>SYAVRDALASIAAIPKIEVHISNVHTREEFRHTSVTVPVCQGEIDGAGLGGYVSAML<br>TLVEMTKSN**GSWGSLEHHHHHH** |
| I3-B3 | *MTDEPVATEEPAARDEVRRAEALGGQAAEQLA*MKILVINGPNINFLGIREKGIYGP<br>FNYDFLVELINKTAELLNVKVEVFQSNHEGAIIDKLQEAYYNDVDGIVINPGAFTHY<br>SYAVRDALASIAAIPKIEVHISNVHTREEFRHTSVTVPVCQGEVDGAGLGGYVSAM<br>ERLVSMTKSN**GSWGSLEHHHHHH** |
| I3-B5 | *MTDEPVATEEPAARDEVRRAEALGGQAAEQLA*MKILVINGPNINFLGIREKGIYGP<br>FNYDDLVDLINKTAEFLNVKVEVFQSNHEGAIIDKLQEAYYNDVDGIVINPGAFTHY<br>SYAVRDALASIAAIPKIEVHISNVHTREEFRHTSVTVPVCQGQVDGAGLAGYVQAM<br>FELTSMTKSN**GSWGSLEHHHHHH** |
| I3-D12 | MKAFEFLYEDFQRGLTVVLDKGLPPKFVEDYLKVCGDYIDFVKFGWGTSAVIDRD<br>VVKEKINYYKDWGIKVYPGGTLFEYAYSKGEAAEFIAECKKLGFEAVEISDGSSDIS<br>LDDRKAAIWGAKWAGFMVLTEVGKKMPDKDKQLTIDDRIKLINFDLDAGADYVIIE<br>GRESGKGIGLFDKEGKVKENELDVLAKNVDINKVIFEAPQKSQQVAFILKFGSSVN<br>LANIAFDEVISLETLRRGLRGDTFGKV**LEHHHHHH** |
| I3-A7<br>(bacterial expression) | MGAIVVTGIPGVGKTTVMQKAAEGSPLPRVPLEGVMYGVAKRMGLVKDIDEMRR<br>LSPDVQKEVQKKAAERIAALGDVILDTHCTIKTPKGYLPGLPRWVLEKLRPSVILLV<br>EADPKEIYGRRLKDETRNRDPDSEEEIAEHQMMNRAAAMAYASLSGATVKIVFNH<br>DNRLDDAVRDAAPVLGN**LEHHHHHH** |
| I3-A7<br>(mammalian expression) | (MDSKGSSQKGSRLLLLLVVSNLLLPQGVLA)MGAIVVTGIPGVGKTTVMQKAAEG<br>SPLPRVPLEGVMYGVAKRMGLVKDIDEMRRLSPDVQKEVQKKAAERIAALGDVIL<br>DTHCTIKTPKGYLPGLPRWVLEKLRPSVILLVEADPKEIYGRRLKDETRNRDPDSE<br>EEIAEHQMMNRAAAMAYASLSGATVKIVFNHDNRLDDAVRDAAPVLGN**GSHHHH<br>HH** |
| I3-A7cp | (MWWRLWWLLLLLLLLWPMVWAAA)**HHHHHHGSGSGS**SEEEIEEHQLINRYAAM<br>AYAVLSGATVKIVFNHDNRLDDAVRDAAPVLHEGEGVIVVTGIPGVGKTTVMQKAA<br>EGSPLPRVPLEGVMYGVAKRMGLVKDIDEMRRLSPDVQKEVQKKAAERIAALGD<br>VILDTHCTIKTPKGYLPGLPRWVLEKLRPSVILLVEADPKEIYGRRLKDETRNRDPD |
| I3-A7cp-N46-N62 | (MWWRLWWLLLLLLLLWPMVWAAA)**HHHHHHGSGSGS**SEEEIEEHQLINRYAAM<br>AYAVLSGATVKIVFNHNNTLDDAVRDAAPVLHNGTGVIVVTGIPGVGKTTVMQKAA<br>EGSPLPRVPLEGVMYGVAKRMGLVKDIDEMRRLSPDVQKEVQKKAAERIAALGD<br>VILDTHCTIKTPKGYLPGLPRWVLEKLRPSVILLVEADPKEIYGRRLKDETRNRDPD |

| | |
|---|---|
| I3-A7cp-N111 | (MWWRLWWLLLLLLLLWPMVWAAA)**HHHHHHGSGSGS**SEEEIEEHQLINRYAAM<br>AYAVLSGATVKIVFNHDNRLDDAVRDAAPVLHEGEGVIVVTGIPGVGKTTVMQKAA<br>EGSPLPRVPLEGVMYGVAKRMGLVKNITEMRRLSPDVQKEVQKKAAERIAALGD<br>VILDTHCTIKTPKGYLPGLPRWVLEKLRPSVILLVEADPKEIYGRRLKDETRNRDPD |
| I3-A7cp-N46-N62-NonAssembling | (MWWRLWWLLLLLLLLWPMVWAAA)**HHHHHHGSGSGS**SEEEIEEHQLINRYAAM<br>AYAVLSGATVKIVFNHNNTLDDAVRDAAPVLHNGTGVIVVTGIPGVGKTTVMQKAA<br>EGMNIEFVTYGTEMFEVAKSMGLVKDRDEMRRLSPDVQKEVQKKAAENIAAKGN<br>VILDTHCTIKTPKGYLPGLPRWVLEKLRPSVILLVEADPKEIYGRRLKDETRNRDPD |
| I3-A7cp-N62 | (MWWRLWWLLLLLLLLWPMVWAAA)**HHHHHHGSGSGS**SEEEIEEHQLINRYAAM<br>AYAVLSGATVKIVFNHDNRLDDAVRDAAPVLHNGTGVIVVTGIPGVGKTTVMQKAA<br>EGSPLPRVPLEGVMYGVAKRMGLVKDIDEMRRLSPDVQKEVQKKAAERIAALGD<br>VILDTHCTIKTPKGYLPGLPRWVLEKLRPSVILLVEADPKEIYGRRLKDETRNRDPD |
| TH-I3-A7 | (MDSKGSSQKGSRLLLLLVVSNLLLPQGVLA)VAPLHLGKCNIAGWILGNPECESLS<br>TASSWSYIVETSNSDNGTCFPGNFINYEELRCQLSSVSSFERFEIFPKTSSWPNH<br>DSNKGVTAACPHAGAKSFYKNLIWLVKKGNSYPKLNQSYINDKGKEVLVLWGIHH<br>PSTTADQQSLYQNEDTYVFVSTSRYDKVFKPIIATRPKVRDQEGRMNYYWTLVEP<br>GDKITFEATGNLVVPRYAFTMERNAGSGSGSCIENINSKIYHIENKLALDRLILALEA<br>KEAGKDWEGIEKVIKELEKVAKDDEEAKKFLEKKKKELEEEKEEWKREEEIEEHQ<br>LINRYAAMAYAVLSGAEVKIVFNHDNRLDDAVRDAAPVLHNGTGVIVVTGIPGVGK<br>TTVMQKAAEGSPLPRVPLEGVMYGVAKRMGLVKDIDEMRRLSPDVQKEVQKKAA<br>ERIAALGDVILDTHCTIKTPKGYLPGLPRWVLEKLRPSVILLVEADPKEIYGRRLKD<br>ETRNRDPD**GSGSHHHHHH** |
| TH-I53_dn5<br>(A component : B component) | MGKYDGSKLRIGILHARGNAEIILELVLGALKRLQEFGVKRENIIIETVPGSFELPYG<br>SKLFVEKQKRLGKPLDAIIPIGVLIRGSTAHFDYIADSTTHQLMKLNFELGIPVIFGVL<br>TTESDEQAEERAGTKAGNHGEDWGAAAVEMATKFN**HHHHHH** :<br>(MDSKGSSQKGSRLLLLLVVSNLLLPQGVLA)VAPLHLGKCNIAGWILGNPECESLS<br>TASSWSYIVETSNSDNGTCFPGNFINYEELRCQLSSVSSFERFEIFPKTSSWPNH<br>DSNKGVTAACPHAGAKSFYKNLIWLVKKGNSYPKLNQSYINDKGKEVLVLWGIHH<br>PSTTADQQSLYQNEDTYVFVSTSRYDKVFKPIIATRPKVRDQEGRMNYYWTLVEP<br>GDKITFEATGNLVVPRYAFTMERNAGSGSGSCIEHIENEIAELAYLLGELAYKLGEY<br>RIAIRAYRIALKSDPNNAEAWYNLGNAYYKQGRYREAIEYYQKALELDPNNAEAW<br>YNLGNAYYERGEYEEAIEYYRKALRLDPNNADAMQNLLNAKMREEGGWELQ**HH**<br>**HHHH** |
| Full-length HA ectodomain trimer (A/Michigan/45/2015), including stabilizing mutations from Milder et al. (1) | (MDSKGSSQKGSRLLLLLVVSNLLLPQGVLA)DTLCIGYHANNSTDTVDTVLEKNV<br>TVTHSVNLLEDKHNGKLCKLRGVAPLHLGKCNIAGWILGNPECESLSTASSWSYIV<br>ETSNSDNGTCFPGDFINYEELREQLSSVSSFERFEIFPKTSSWPNHDSNKGVTAA<br>CPHAGAKSFYKNLIWLVKKGNSYPKLNQSYINDKGKEVLVLWGIHHPSTTADQQS<br>LYQNADAYVFVGTSRYSKKFKPEIATRPKVRDQEGRMNYYWTLVEPGDKITFEAT<br>GNLVVPRYAFTMERNAGSGIIISDTPVHDCNTTCQTPEGAINTSLPFQNIHPITIGK<br>CPKYVKSTKLRLATGLRNVPSIQSRGLFGAIAGFIEGGWTGMVDGWYGYHWQNE<br>QGSGYAADLKSTQNAIDKITNIVNSVIEKMNTQFTAVGKEFNHLEKRIENLNKKVDD<br>GFLDIWTYNAELLVLLINERTLDYHDSNVKNLYEKVRNQLKNNAKEIGNGCFEFYH<br>KCDNTCMESVKNGTYDYPKYSEEAKLNREKIDGVGSGYIPEAPRDGQAYVRKDG<br>EWVLLSTFLGSGLNDIFEAQKIEWHEG**HHHHHH** |

Signal peptides are indicated in parentheses and purification tags are bolded and underlined.
Italicized and underlined sections of sequences from I3-A6, I3-B2, I3-B3, I3-B5 represent mistakenly added sequence from an unrelated thermophilic organism, *Thermasporomyces composti*, due to an error in BLAST retrieval of full length genomic sequences. This region was not resolved by cyro-EM and is unlikely to have an impact on nanoparticle assembly.

| Table S3. Cryo-EM Data Collection Statistics | | | |
|---|---|---|---|
| | I3-A6<br>PDB: 9CLZ<br>EMD: 45734 | I3-D12<br>PDB: 9CM1<br>EMD: 45736 | I3-A7<br>PDB: 9CM0<br>EMD: 45735 |
| **Data collection and processing** | | | |
| Magnification (×) | 105,000 | 105,000 | 105,000 |
| Voltage (kV) | 300 | 300 | 300 |
| Electron exposure ($e^-/Å^2$) | 52 | 52 | 52 |
| Defocus range (μm) | -0.5 to -2.0 | -0.5 to -2.0 | -0.5 to -2.0 |
| Pixel Size (Å) | 0.843 | 0.843 | 0.843 |
| Symmetry imposed | I | I | I |
| Final particle images (no.) | 290,122 | 321,856 | 165,931 |
| Map resolution (Å) | 2.5 | 2.3 | 3.5 |
| FSC threshold | 0.143 | 0.143 | 0.143 |
| **Validation** | | | |
| MolProbity score | 1.03 | 0.53 | 0.57 |
| Clashscore | 0.26 | 0.08 | 0.18 |
| Poor rotamers (%) | 1.94 | 0.33 | 0.7 |
| Ramachandran plot | | | |
| Favored (%) | 96.68 | 99.15 | 97.07 |
| Allowed (%) | 100 | 100 | 100 |
| Disallowed (%) | 0 | 0 | 0 |
